# TAp73 regulates mitochondrial dynamics and multiciliated cell homeostasis through an OPA1 axis

**DOI:** 10.1101/2023.03.23.533672

**Authors:** Niall Buckley, Andrew Craxton, Xiao-Ming Sun, Emanuele Panatta, Lucia Pinon, Jaime Llodrá, Nobuhiro Morone, Ivano Amelio, Gerry Melino, L. Miguel Martins, Marion MacFarlane

**Affiliations:** MRC Toxicology Unit, University of Cambridge, Gleeson Building, Tennis Court Road, Cambridge CB2 1QR, UK; Department of Experimental Medicine, University of Rome "Tor Vergata", 00133 Rome, Italy; Division for Systems Toxicology, Department of Biology, University of Konstanz, Konstanz, Germany

**Keywords:** p53 family, mitochondria, ciliogenesis, COPD

## Abstract

Dysregulated mitochondrial fusion and fission has been implicated in the pathogenesis of numerous diseases. We have identified a novel function of the p53 family protein TAp73 in regulating mitochondrial dynamics. TAp73 regulates the expression of Optic atrophy 1, a protein responsible for controlling mitochondrial fusion, cristae biogenesis and electron transport chain function. Disruption of this axis results in a fragmented mitochondrial network and an impaired capacity for energy production *via* oxidative phosphorylation. Owing to the role of OPA1 in modulating cytochrome *c* release, TAp73^-/-^ cells also display an increased sensitivity to apoptotic cell death, e.g., *via* BH3-mimetics. We also show that the TAp73/OPA1 axis has functional relevance in the upper airway, where TAp73 expression is essential for multiciliated cell differentiation and function. Consistently, ciliated epithelial cells of *Trp73^-/-^* (global p73 KO) mice display decreased expression of OPA1 and perturbations of the mitochondrial network, which may drive multiciliated cell loss. In support of this, *Trp73* and *OPA1* gene expression is decreased in COPD patients, a disease characterised by alterations in mitochondrial dynamics. We therefore highlight a potential mechanism involving the loss of p73 in COPD pathogenesis. This work also adds to the growing body of evidence for growth-promoting roles of TAp73 isoforms.

## INTRODUCTION

The transcription factor p73, together with p53 and p63, is a member of the p53 family (1–3). On account of a common ancestor and a highly conserved DNA-binding domain, each family member has the ability to activate a shared set of genes responsible for cell-cycle arrest and apoptosis following DNA damage (4–6). However, they also have distinct mechanisms of regulation by upstream signalling pathways and post-translational modifications (7,8). The *Trp73* gene consists of two alternative promoters, giving rise to transcriptionally proficient transactivation (TAp73) and anti-apoptotic N-terminally deleted dominant negative (ΔNp73) isoforms (8). Moreover, further complexity is generated through C-terminal alternative splicing, which modulates p73 function (9,10).

In addition to its p53-like functions, p73 fulfils unique DNA-damage independent roles (11). Indeed, the expanding set of TAp73 functions in embryonic development, tissue homeostasis and cancer demonstrate its vast functional pleiotropy (12). The generation of p73 knockout mouse models has uncovered important roles in development of the nervous system (11,13), the control of metabolism (14–16), spermatogenesis (17), angiogenesis (18) and multiciliogenesis (19,20). However, the molecular underpinnings of the diverse tissue dysfunction require further investigation. One such potential mechanism is a role of TAp73 in the control of metabolism and mitochondrial function (14). For example, TAp73 deletion from mouse embryonic fibroblasts (MEFs) leads to a downregulation of the complex IV subunit *Cox4i1*. This results in reduced complex IV activity, and a concomitant decrease in cellular ATP, oxygen consumption, and a premature ageing phenotype *in vivo* (16). Additional molecular mechanisms by which TAp73 regulates mitochondrial function and metabolism are through the direct transcriptional control of *GLS2*, the gene encoding glutaminase type 2 (21), glutamine metabolism (22), and the synthesis of serine (23).

We present data highlighting a novel transcriptional role for TAp73 in regulating the mitochondrial shaping protein Optic Atrophy 1 (OPA1) *in vitro* and *in vivo*. Functionally, OPA1-dependent mitochondrial fusion supports increased rates of mitochondrial oxidative phosphorylation (24). Moreover, independently of fusion, OPA1 also regulates respiratory complex assembly and OXPHOS efficiency, owing to its function in maintaining cristae morphogenesis (25,26). The importance of OPA1 in the maintenance of mitochondrial structure, genome, and function is also evident from mouse and patient models where mutation or loss of OPA1 gives rise to several pathophysiological outcomes including wasting of skeletal muscle and autosomal dominant optic atrophy (27,28). We show that ablation of TAp73 *in-vitro* elicited a downregulation of OPA1 expression, a concomitant fragmentation of the mitochondrial network, impaired bioenergetic function, and increased sensitivity to apoptosis. We therefore present an overarching mechanism by which TAp73 regulates mitochondrial dynamics and respiratory function.

We have also investigated the TAp73/OPA1 axis *in vivo*, focussing on the airway ciliated epithelium in *Trp73^-/-^* mice, which have a global deletion of all p73 isoforms. Importantly, although the generation of motile cilia is severely abrogated in both *Trp73* and TAp73 knockout mice, TAp73 isoforms are necessary and sufficient for functional multiciliogenesis (19). Multiciliated cells lacking TAp73 are characterised by short appendages with impaired mucociliary clearance. Multiciliogenesis is a highly energy dependent process and is intricately linked to mitochondrial function (29). Integral to cilia microtubule polymerisation is the interaction of ATP at the exchangeable GTP site of tubulin (30), with insufficient ATP generation leading to decreased microtubule stability (31). Such a phenomenon was evident in OPA1- deficient neutrophils, which produce insufficient ATP for neutrophil extracellular trap formation (32). As such, the regulation of OPA1 by TAp73 in the ciliated epithelium suggests that TAp73-dependent metabolic regulation may participate in the multiciliogenesis process (14). In support of this, we show that *Trp73* ablation led to a decrease in OPA1 expression and altered mitochondrial morphology in the ciliated epithelium of the mouse airway. This therefore represents a potential mechanism underpinning multiciliated cell loss in *Trp73* null mice and may also be relevant in chronic obstructive pulmonary disease (COPD) pathogenesis, as patient cohorts display decreased *Trp73* and *OPA1* expression. Furthermore, this novel function also represents a growth-promoting function of TAp73 isoforms. Indeed, although p53 is lost or mutated in about half of human cancers, the same is not true for p73 (33,34). On the contrary, certain p73 isoforms are overexpressed in a range of cancers and influences disease prognosis (35–38). This work therefore further underlines the highly diverse role TAp73 plays in tumorigenesis.

## RESULTS

### TAp73 regulates OPA1 expression and binds with the promoter region

TAp73 has a number of roles in regulating cellular metabolism. We have previously reported a correlation between TAp73 and OPA1 expression in H1299 cells, suggesting that the control of metabolic processes by TAp73 might also occur through the regulation of mitochondrial morphology (9). To address this possibility, genome-wide binding sites for TAp73α, TAp73β and p53 were interrogated from previously published ChIP-seq data (39). The binding profile upstream of the OPA1 transcription start site (TSS) showed an enrichment of reads at distinct loci in immunoprecipitated samples, indicating TAp73 binding (figure 1a). Moreover, this enrichment was evident for both TAp73α and TAp73β isoforms, suggesting that the SAM domain of TAp73α is not required for this interaction. Furthermore, p53 binding was evident adjacent to the OPA1 transcription start site, suggesting that the ability of TAp73 to bind the OPA1 promoter region may be conserved throughout the p53 family (figure 1a, S1a). The data sets also provided the opportunity to explore the possibility that TAp73 may regulate additional genes that influence mitochondrial dynamics. In support of this, defined peaks were evident for TAp73α, TAp73β and p53 at the *MFN-2*, *MFF* and *FIS1* gene promoter regions (figure S1a). In addition, GO enrichment analysis of 1769 TAp73 target genes identified using *in-situ* mouse tracheal ChIP-seq displayed an overrepresentation of genes that regulate mitochondrial membrane organisation (Gene Ontology ID: 0007006). These included *OPA1*, *MFN2* and *PPARGC1A*, the gene encoding a subunit of PGC-1α (figure S1b).

**Figure 1.**
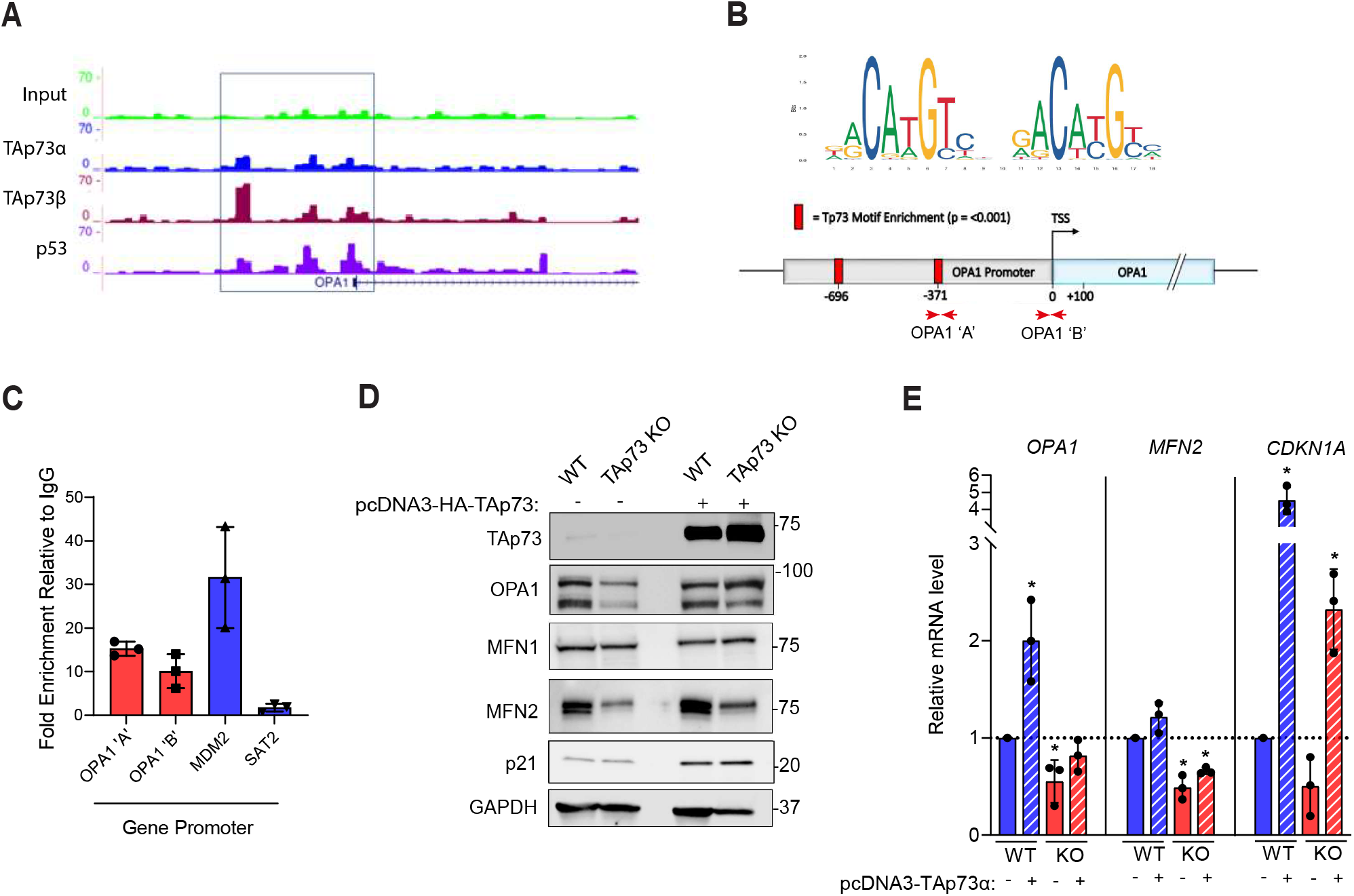
TAp73 regulates the expression of OPA1. (**A**) Interrogation of ChIP-seq data indicated binding of TAp73α, TAp73β and p53 to the putative OPA1 promoter region. Sequencing read files were obtained from the GEO data set GSE15780, and tracks shown for the indicated transcription factors at selected genes. (**B-C**) Targeted ChIP of TAp73 bound chromatin. RT-qPCR primers were designed in the promoter region of the OPA1 gene (OPA1 ‘A’ and OPA1 ‘B’). Red squares indicate regions enriched for the Trp73 motif (p<0.001). qPCR was performed to quantify the fold enrichment of the OPA1 promoter region in IP sample relative to IgG control. Enrichment of MDM2 and SAT2 promoter regions was assayed as positive and negative controls, respectively. qPCR was carried out on 3 independent ChIP experiments and data shown as individual data points ± SD (n=3). (**D**) Representative Western blot of mitochondrial fusion proteins in TAp73 KO and WT control. Cells were transfected with either EV or TAp73α expression construct for 24h. (**E**) RT-qPCR was performed against *OPA1*, *MFN2* and *CDKN1A* genes and expression values calculated using the ΔΔCt method, relative to WT empty vector control. Data shown as mean ± SD (n=3). (*) P ≤ 0.05 (Student’s t-test, comparison of indicated condition with WT EV control).

We subsequently selected the OPA1 gene for further investigation due to its important roles in both mitochondrial fusion, cristae morphogenesis and respiratory function. Targeted IP of TAp73-bound chromatin followed by qPCR of the OPA1 promoter showed an enrichment at two regions upstream of the OPA1 TSS, relative to IgG control (figure 1b,c). This therefore validated the ability of overexpressed TAp73α to bind the OPA1 promoter. Moreover, the OPA1 promoter region also showed a significant enrichment for the *Trp73* binding motif at two loci (figure 1b).

Next, we generated TAp73 knockout (KO) H1299 cell lines using CRISPR/Cas9 targeting to investigate the effect of TAp73 ablation OPA1 expression (figure S2). Genetic ablation of TAp73 isoforms resulted in a decrease in OPA1 expression at the protein and mRNA level (figure 1d,e). Moreover, ectopic expression of TAp73 was sufficient to restore OPA1 expression in TAp73 KO cells. Together, this places OPA1 downstream of TAp73 and suggests such an axis occurs *via* transcriptional regulation. We also observed a similar pattern of expression of MFN2, which had downregulated expression in TAp73 KO cells (figure 1d,e). Together with ChIP-seq data (figure S1a), this suggests *MFN2* may also be a TAp73 target gene, albeit its expression was not rescued by ectopic expression of TAp73.

### Disruption of the TAp73/OPA1 axis alters mitochondrial morphology and bioenergetics

Following the identification of an axis between TAp73 and OPA1, we investigated the morphology and function of the mitochondrial network in TAp73 KO cells. First, we addressed whether the observed OPA1 depletion was sufficient to impair mitochondrial fusion and alter steady-state mitochondrial dynamics. Immunofluorescence staining for ATP5B indicated a striking fragmentation of the mitochondrial network in TAp73 KO cells compared with WT control cells (figure 2a). This was also recapitulated by siRNA knockdown of TAp73 (Figure S3). Mitochondrial fragmentation was also quantified using the Intellesis trainable segmentation module (Zeiss), which quantified mean mitochondrial area across biological replicates (figure 2c). To confirm the observed mitochondrial phenotype was driven by TAp73 knockout, ectopic expression of TAp73 rescued a fused mitochondrial state. Likewise, mitochondrial fragmentation was rescued with ectopic OPA1 expression (figure 2a-c), placing OPA1 downstream of TAp73. However, the OPA1 rescue was not as effective as observed following transfection with TAp73 expression vector, suggesting the activity of additional regulators of fusion and fission may remain deficient. We also observed perturbations in mitochondrial morphology in TEM micrographs of TAp73 KO cells (figure 2d,e). Overall, these data demonstrate a role for TAp73 in regulating mitochondrial dynamics.

**Figure 2.**
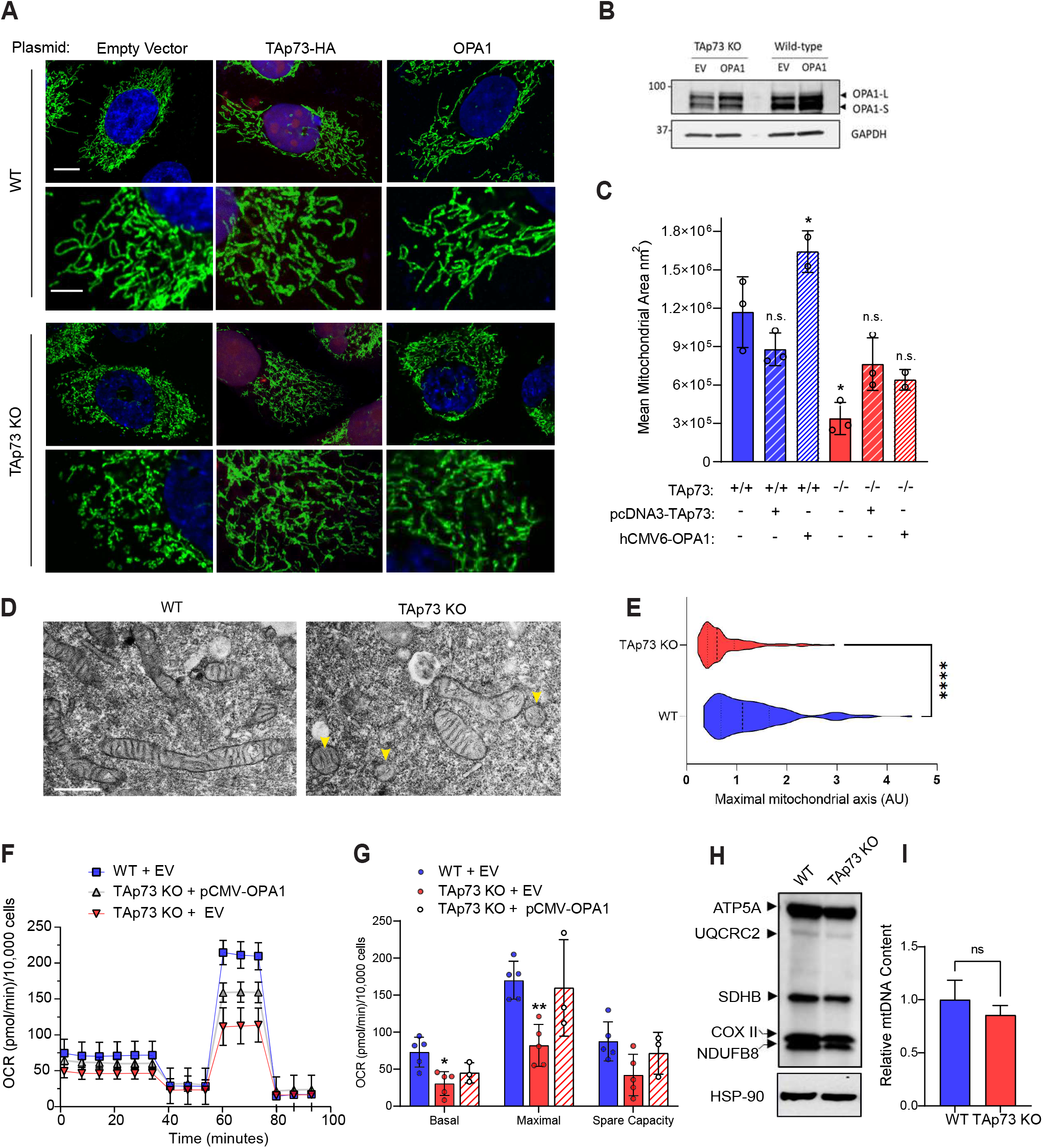
TAp73 KO cells display fragmented mitochondria and impaired ETC function. (A) WT or TAp73 KO H1299 cells were transfected with the indicated plasmids for 24h and IF carried out against ATP5B (green) with DAPI nuclear counterstain (blue). Cells transfected with HA-TAp73α expression plasmid were stained for HA as a transfection control (red). Scale bar = 10 μm. (B) Representative western blot of OPA1 expression following transfection of WT and TAp73 KO cells with pCMW-OPA1 construct. (C) Quantification of mitochondrial morphology from (a) using Zeiss Intellesis module, trained to segment individual mitochondria. Statistical significance compared to WT EV control was calculated using Student’s T-test; (*) p < 0.1, (ns) not significant (n=3). (D) Transmission electron micrographs of mitochondrial morphology from WT and TAp73 KO cells. Scale bar = 100 nm. (E) Mitochondrial length measurements obtained from (D). (****) p<0.0001 in Student’s t-test. A minimum of 100 mitochondria were measured from n=3 independent biological replicates. (**F-G**) Mitochondrial stress test performed on Seahorse XFe96 analyser. Canonical mitochondrial inhibitors injected sequentially as labelled (Oligomycin = 2μM, FCCP = 500nM, Antimycin A/ Rotenone = 2μM). The indicated mitochondrial stress test parameters were calculated from OCR data. Data were corrected for non-mitochondrial OCR, normalised to cell number, and are shown as mean ± SD (n=3). (*) P ≤ 0.05 and (**) P ≤ 0.01 in Student’s T-test relative to WT control. (H) Western blot of the indicated ETC subunits in Wild-type and TAp73 KO cells, obtained using OXPHOS antibody cocktail. (I) qPCR against *mt-CO2*, expressed relative to expression of nuclear encoded β*2- microglobulin*. Relative expression was calculated using the ΔΔCt method and expressed as a percentage of wild-type control (n=2). n.s = not significant in Student’s t-test.

We next investigated the impact of TAp73 knockout and the resultant changes in mitochondrial morphology on mitochondrial bioenergetic function. Analysis using the Seahorse extracellular flux assay revealed that TAp73 knockout cells displayed a decrease in basal OCR, maximal OCR and spare capacity, which was rescued upon overexpression of OPA1 (figure 2f,g). These data therefore indicated that disruption of the TAp73/OPA1 axis elicited an impairment in ETC function, probably due to the role of OPA1 in maintaining cristae architecture and respiratory complex efficiency. In support of this theory, we also found a disruption in cristae architecture in TAp73 KO cells (figure 2c-e). Moreover, we were able to eliminate a depletion of mtDNA and/or a decrease in translation of ETC complexes as a mechanism driving the observed impairment in oxidative phosphorylation, as there was not a significant decrease in either species (figure 2h,i).

### TAp73 knockout cells display an increased sensitivity to apoptosis induction with BH3-mimetics

The functions of OPA1 in regulating mitochondrial fusion and cristae remodelling are inextricably linked with the execution of apoptosis. Upstream of caspase-9 activation, mitochondria fragment to facilitate the release of cytochrome *c* from cristae into the cytosol where if binds with APAF1 and activates the assembly of the apoptosome (40). As such, blocking of mitochondrial fission inhibits cytochrome *c* release and cell death. Concurrently, overexpression of OPA1 inhibits mitochondrial fission and apoptotic cristae remodelling, thereby rendering cells resistant to apoptosis induction by the intrinsic pathway (26).

To interrogate the impact of altered mitochondrial morphogenesis on apoptosis induction in TAp73 KO cells, mitochondrial outer membrane permabilisation was triggered by treating cells with BH3-mimetics (figure 3a). This approach allowed us to investigate a mitochondrial vulnerability in TAp73 KO cells whilst circumventing the role of TAp73 in orchestrating upstream modulators of apoptosis, such as PUMA and NOXA (41). Due to the upregulation of multiple members of the pro-survival Bcl-2 family of proteins in H1299 cells, double agent treatment with ABT-737 (Bcl-2 and Bcl-xL inhibitor) and S63845 (Mcl-1 inhibitor) was necessary to induce apoptosis (42). Quantification of the kinetics of apoptosis induction indicated that TAp73 KO cells were more sensitive to 0.5 μM and 1 μM combination treatment of BH3-mimetic (figure 3a). Importantly, the observed sensitivity did not appear to be mainly driven by a downregulation of the pro-survival proteins targeted by BH3-mimetics (figure 3b). It was also evident in untreated cells that TAp73 knockout induced alterations in cristae architecture (figure 3c). The width of mitochondrial cristae was significantly increased in KO cells, whilst the cristae density was profoundly decreased (figure 3e,d). This finding was consistent with a decrease in OPA1 expression, and probably drives an increased propensity for apoptosis.

**Figure 3.**
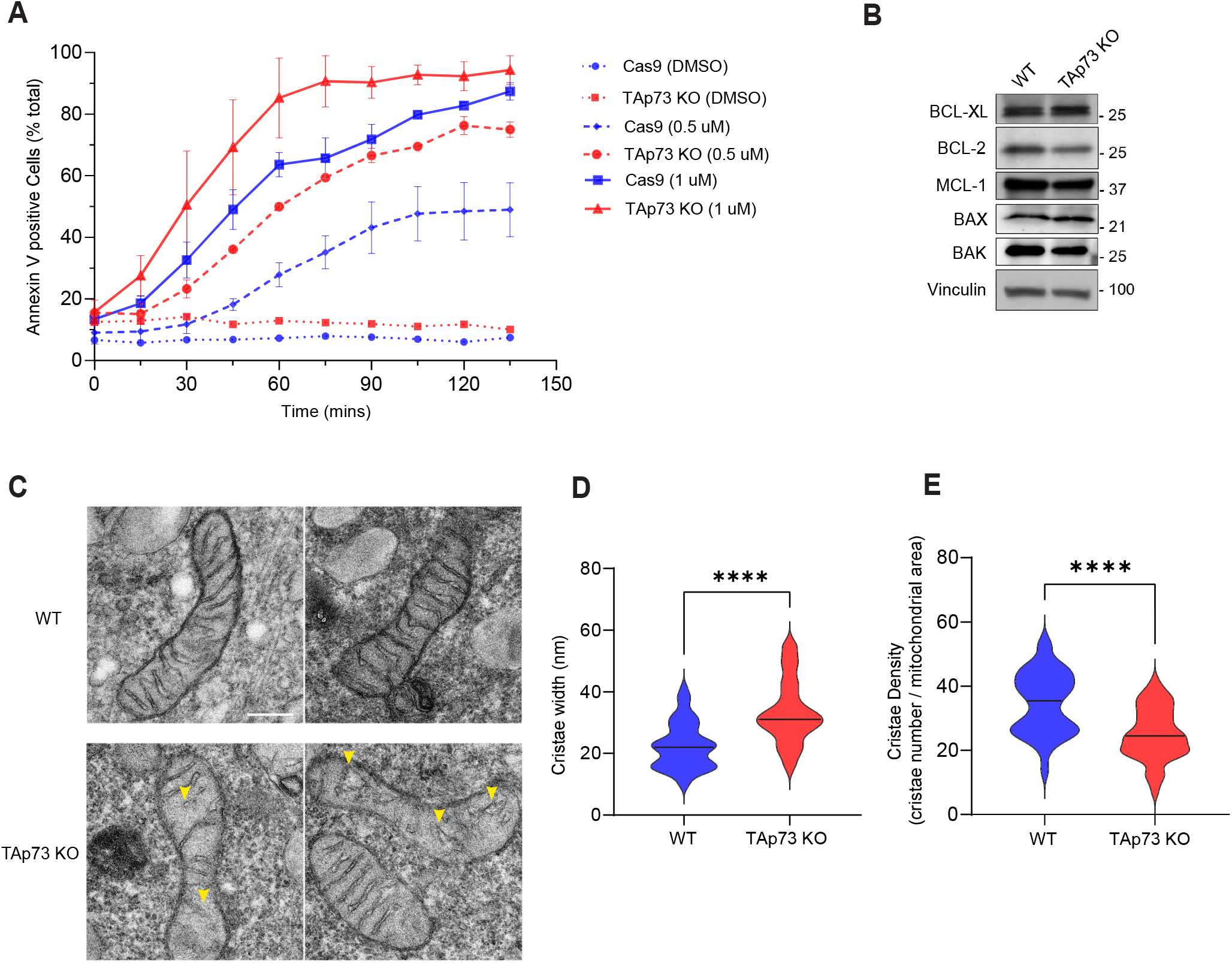
TAp73 KO cells are sensitised to apoptosis induced by BH3-mimetics. (A) Kinetics of apoptosis induction was tracked using AnnexinV/APC dye following treatment with BH3-mimetic. Wild-type or TAp73 KO H1299 cells were treated with combination treatment of ABT-737 and S63845 at the indicated concentrations. Data points plotted as mean ± SD from n=6 technical replicates. (B) Representative Western blot against pro-apoptotic and anti-apoptotic proteins of the Bcl-2 family in WT and TAp73 KO cell lines. (C) Representative TEM micrographs of mitochondrial ultrastructure in WT and TAp73 KO cells. Yellow arrows highlight regions of disorganisation or loss of cristae in TAp73 KO cells. Scale bar = 200 nm. (**D-E**) Quantification of mitochondrial cristae width (d) and cristae density (e) in TEM micrographs obtained from WT and TAp73 KO H1299 cells. Measurements were obtained from two biological replicates and (****) p<0.0001 in Student’s t-test.

### The TAp73/OPA1 axis is functionally relevant in the airway ciliated epithelium

Our finding that an axis exists between TAp73 and OPA1 that regulates mitochondrial dynamics prompted us to address the biological significance of this relationship *in vivo*. To do so, we analysed epithelial cells of the upper airway in *Trp73^-/-^* mice, a tissue ordinarily with high expression of TAp73 isoforms in mouse and human (43,44). Specifically, TAp73 plays an important role in multiciliated cells (MCCs), where it has been shown to regulate the expression of key genes responsible for multiciliated cell function and homeostasis (19,20).

Interestingly, although we observed a structural perturbation of cilia in *Trp73* null mice (figure 4a), the frequency of multiciliated cell loss was not as profound as previously reported (20), as 17% of cells successfully nucleated motile cilia; albeit with a defective architecture. This suggests that TAp73 ablation manifests as a late-stage defect in the multiciliogenesis process, supported by the observation that TAp73 is not expressed in the p63+ progenitor cell compartment (figure S4a). We also observed the presence of FOXJ1 positive cells in the *Trp73^-/-^* epithelium (figure S4b). Together, these findings indicate that additional molecular mechanisms - extending beyond TAp73 mediated fate determination and axonemal gene regulation - may underpin the multiciliated cell phenotype of *Trp73^-/-^* mice.

**Figure 4.**
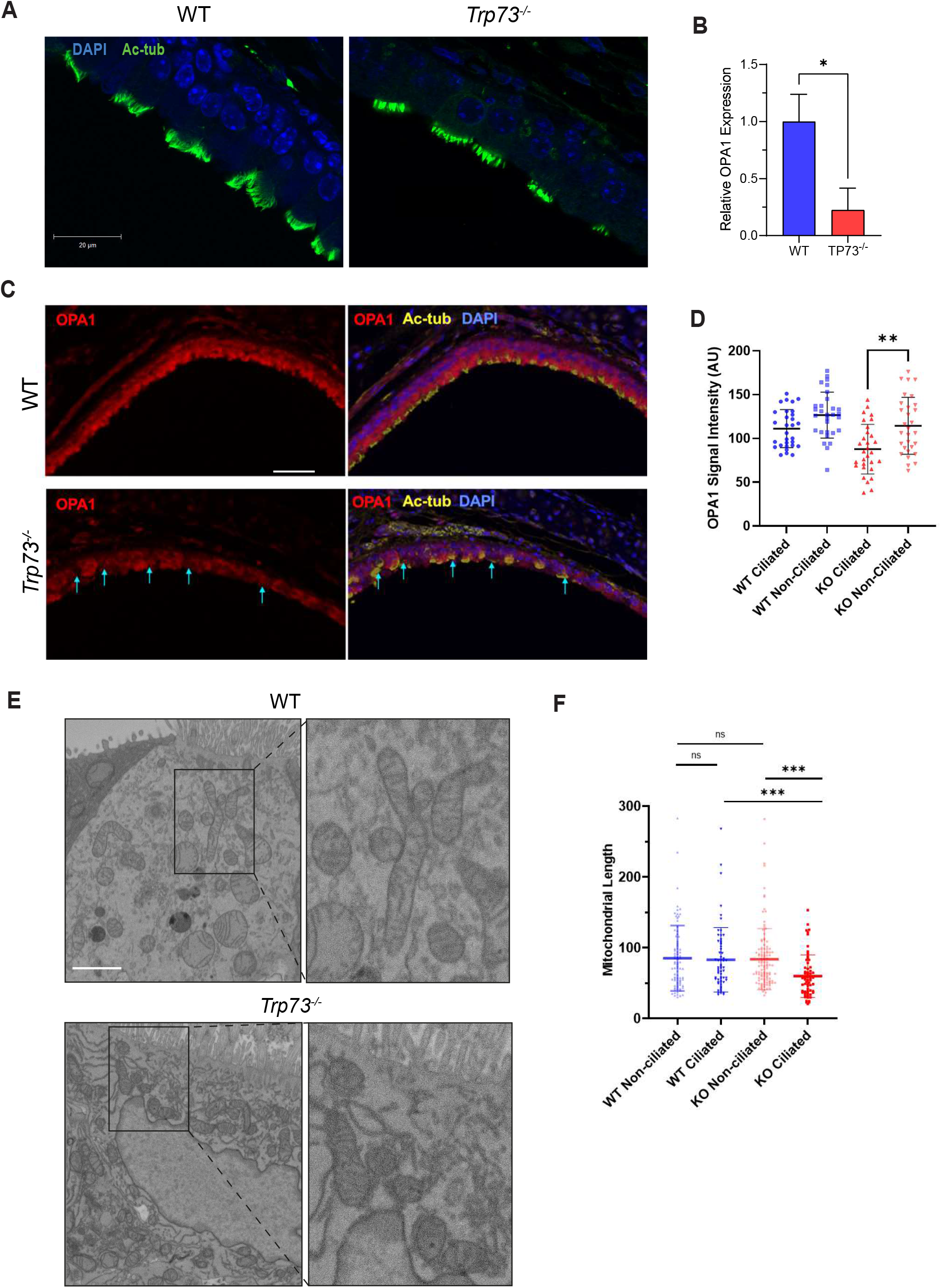
*Trp73^-/-^* mice exhibit decreased OPA1 expression and altered mitochondrial dynamics in the airway ciliated epithelium. (A) Immunofluorescence staining performed against the cilia marker Ac-α-tubulin (green) with DAPI nuclear stain in tracheal cross-sections from WT and *Trp73^-/-^* mice (blue). Scale bar = 20 μm. (B) RT-qPCR for *OPA1* mRNA expression in dissociated tracheal epithelial cells. Mouse trachea (n=3 of each WT and *Trp73^-/-^*) were pooled together to obtain a sufficient number of cells. (C) Multiplexed IHC on mouse trachea against Ac-a-tubulin (yellow), OPA1 (red) and DAPI (blue). Cyan arrows indicate MCCs with low OPA1 expression. (D) Quantification of OPA1 expression in ciliated and non-ciliated cell populations from WT and *Trp73^-/-^* mice. (E) Images of individual sections of MCCs from WT and *Trp73^-/-^* ciliated epithelium obtained by SBF-SEM. Scale bar = 200 nm. (F) Mitochondrial length measurements were obtained from (E), including ciliated and non-ciliated cell populations. n≥100 mitochondria from two samples of each genotype. (***) P ≤ 0.001, n.s = not significant, in t-test.

In support of this hypothesis, multiplexed immunohistochemical (IHC) staining of the *Trp73^-/-^* tracheal epithelium showed a reduction in OPA1 expression (figure 4c,d).

Moreover, co-staining for Acetylated-α-tubulin revealed that OPA1 expression was reduced specifically in the ciliated cell lineage, from which *Trp73* was ablated (cyan arrows). Conversely, non-ciliated cells from *Trp73^-/-^* animals displayed similar OPA1 expression to WT controls (figure 4d). Moreover, RT-qPCR indicated a reduction in OPA1 mRNA in the ciliated epithelium of *Trp73^-/-^* mice, in support of a transcriptional regulation by p73 (figure 4b).

We next addressed whether altered OPA1 expression in ciliated cells of the *Trp73^-/-^* tracheal epithelium drives alterations in mitochondrial morphology and employed SBF-SEM to visualise the mitochondrial network. Individual sections from WT and Trp73^-/-^ ciliated epithelial cells are shown in figure 4e and revealed a fragmented mitochondrial phenotype in knockout mice, which was confined to the MCC lineage (figure 4f), consistent with downregulated OPA1 expression. Furthermore, mitochondria from multiciliated cells were segmented in serial SEM sections to reconstruct the network in 3D (figure S5). Visualisation of the mitochondrial network in tracheal epithelial cells revealed a highly apical localisation of mitochondria in WT animals, consistent with previous reports (45). This pattern of localisation was not observed in *Trp73^-/-^* epithelial cells, suggesting an alteration in mitochondrial trafficking. Overall, these data indicated an alteration in mitochondrial dynamics in the multiciliated cell lineage of *Trp73^-/-^* mice.

### *Trp73* and *OPA1* expression is dysregulated in COPD patients

Chronic obstructive pulmonary disease (COPD) is characterised by a complex disease pathology and driven by multiple underlying mechanisms (46). However, it is often associated with airway multiciliated cell dysfunction, a loss or shortening of motile cilia, and reduced ciliary beating (47,48). Due to this similarity with p73 knockout mouse models, and the requirement for p73 for MCC function, we postulated that alterations in p73 expression could be implicated in COPD pathogenesis. Consistently, there was a highly significant reduction in *Trp73* expression in whole lung homogenates from COPD patients when compared with healthy controls (Lung Genomics Research Consortium; GSE47460) (figure 5a). Furthermore, there was a correlation between *Trp73* and *OPA1* expression across healthy and COPD lung cohort data (figure 5b,c). Indeed, extracting expression data from COPD patients highlighted a clustering of expression profiles with decreased expression of both *Trp73* and *OPA1*, driving a positive Pearson’s coefficient (figure 5d). However, the strength of this relationship may be diminished by the contribution of non-ciliated cell populations, indicated by the highly similar correlation value obtained for *CDKN1A* expression, a known transcriptional target of TAp73 (figure 5e). Overall, these findings highlight a dysregulation of p73 expression in COPD pathogenesis, and correlates with altered OPA1 expression, as observed in the *Trp73^-/-^* mouse airway epithelium (figure 6).

**Figure 5.**
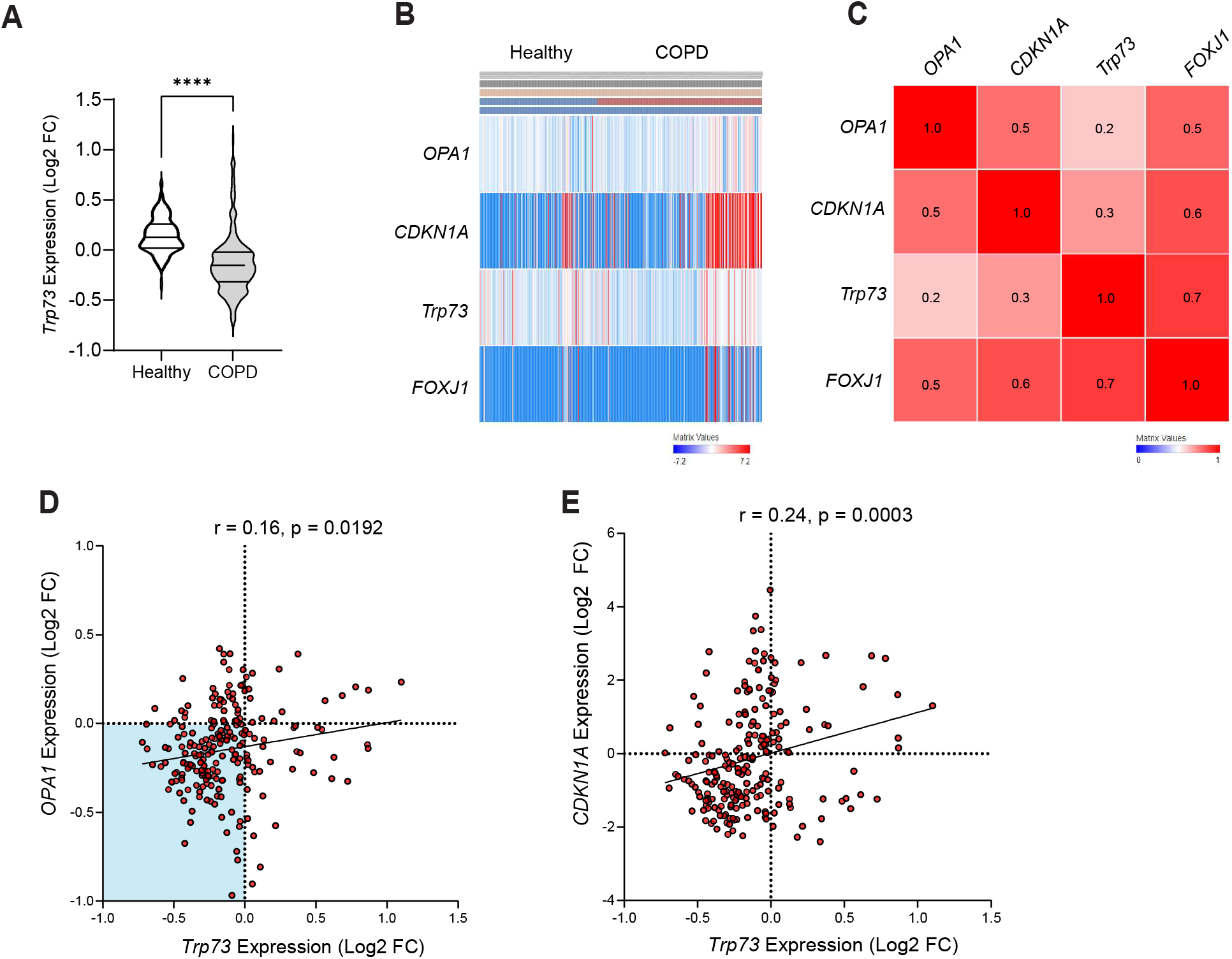
Expression of *Trp73* is decreased in COPD and correlates with *OPA1* expression levels. (A) *Trp73* expression data from healthy and COPD individuals from previously described Lung Genomics Research Consortium (LGRC) cohort (GSE47460; Affymetrix array data from n=157 healthy and n=220 COPD patients). (B) Heatmap of *Trp73, CDKN1A, OPA1* and *FOXJ1* expression in healthy and COPD individuals from the LGRC cohort. The heatmap was generated with the PulmonDB tool using Scipy library in Python, utilising cosine distance and average linkage. Red/blue cells represent positive/negative values. (C) Row similarity matrix indicating the association between each gene across patient data shown in (B). Red shading indicates a positive similarity (measured as 1 - cosine-distance, with similarity values indicated). (**D-E**) Expression data showing the correlation between *Trp73* and *OPA1* (D), or *Trp73* and *CDKN1A* (E) in COPD patients (GSE47460). Expression values are shown as Log2 fold change for the indicated genes relative to healthy control. The strength of the correlation was calculated using Pearson’s coefficient (r).

**Figure 6.**
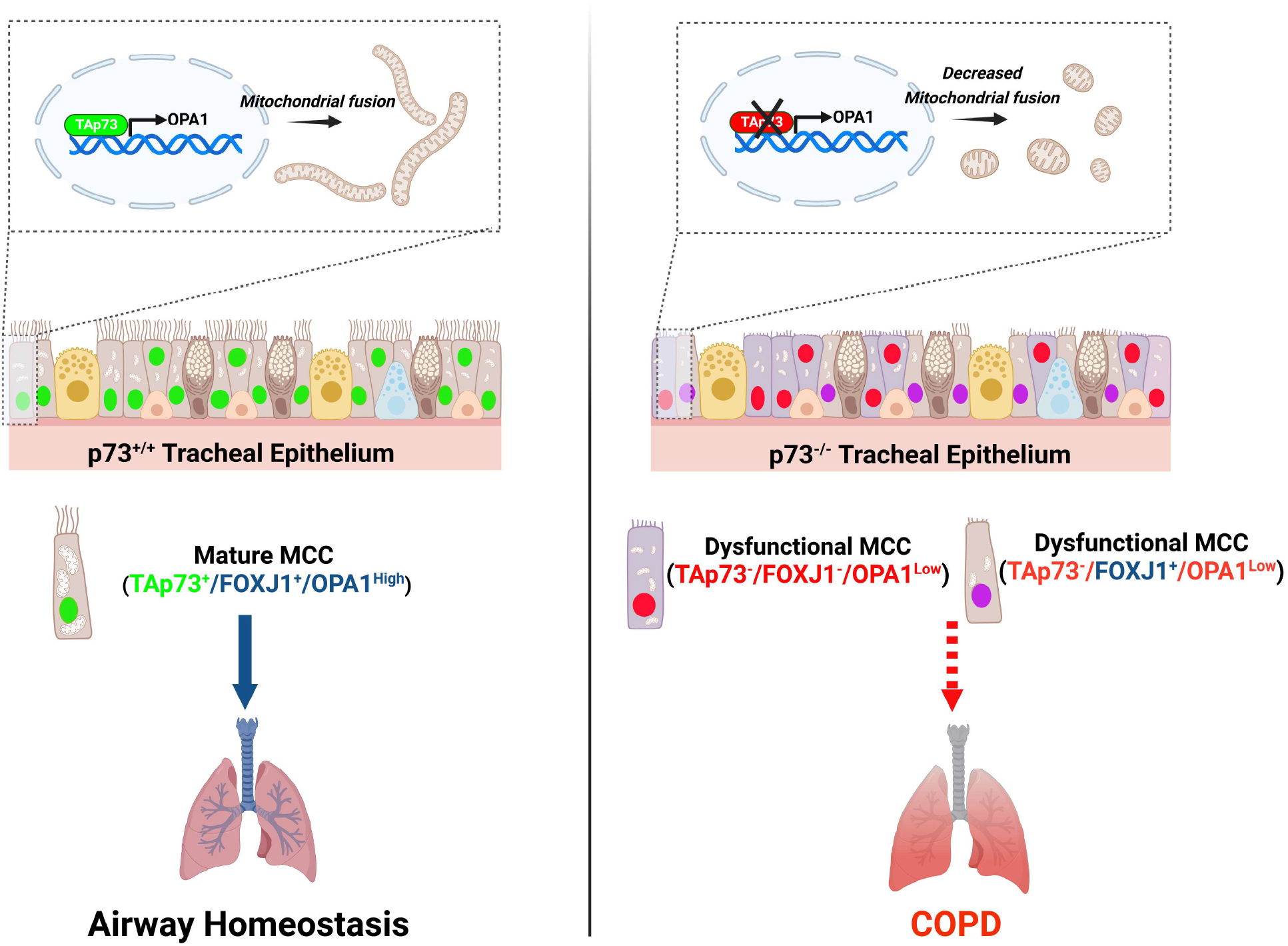
TAp73 regulates mitochondrial dynamics *in vitro* and in airway MCC’s. Schematic illustration showing the identified role of TAp73 in regulating mitochondrial dynamics. TAp73 expression is required for mitochondrial homeostasis *in vitro* and in the ciliated epithelium (green nuclei) (left). Conversely, TAp73 ablation leads to decreased OPA1 expression, mitochondrial fission, and impaired mitochondrial function (right). The *Trp73^-/-^* tracheal epithelium maintains a limited expression of FOXJ1 positive cells (purple), indicating additional mechanisms, such as the observed mitochondrial dysfunction drives MCC loss and COPD pathogenesis.

## DISCUSSION

We have identified a novel role for TAp73 in regulating OPA1 expression and thereby mitochondrial morphology and function. This finding has numerous implications, as the control of mitochondrial dynamics by the p53 family is a key mechanism for the regulation of cell metabolism (including metabolic shifts observed in cancer) (49), tumour cell growth, dissemination(50), and differentiation (51). Indeed, previous evidence has shown that p53 regulates mitochondrial morphology through the transcriptional regulation of MFN2(52). Nonetheless, a role of p73 in regulating fusion and fission genes has not been reported, and may underpin the diverse tissue dysfunction in p73 knockout mouse models (11).

The interrogation of *in-vitro* and *in-vivo* (mouse tracheal) ChIP-seq data sets (20,39) revealed TAp73 isoforms alpha and beta bind to overlapping genomic loci within the promoter region of numerous genes responsible for mitochondrial dynamics, which included OPA1 (figure 1, figure S1). Moreover, highly similar binding patterns were observed for p53, indicative of a conserved ability of the p53 family to regulate mitochondrial shaping proteins. To this end, we performed targeted ChIP which confirmed the ability of TAp73 to bind the OPA1 promoter. The transcriptional regulation of OPA1 by p73 was also interrogated in TAp73^-/-^ H1299 cells, generated by CRISPR/Cas9 targeting (figure 1), and previously reported using an siRNA approach (9). In a CRISPR KO model, the downregulated expression of OPA1 manifested as a profound alteration in mitochondrial morphology. Namely, TAp73 KO cells displayed a fragmented mitochondrial network, and crucially, this was rescued by ectopic expression of OPA1 (figure 2). This therefore demonstrates a clear molecular dependence of the TAp73/OPA1 axis for the maintenance of mitochondrial dynamics (figure 2).

The alterations in OPA1 expression in a TAp73 KO model also demonstrate the importance of intricate control of mitochondrial shaping proteins for functionality of the organelle. OPA1 oligomers perform a myriad of functions that include, but are not limited to, fusion of mitochondria, the modulation of cristae morphogenesis, cytochrome c release, and respiratory complex efficiency (25,53). The status of OPA1 as an effector of these mitochondrial functions therefore adds an additional layer by which TAp73 regulates mitochondrial bioenergetics, as we observed a defect in ETC function that was rescued with ectopic OPA1 expression (figure 2). Moreover, disruption of the TAp73/OPA1 axis appeared to increase sensitivity to apoptosis induced by BH3-mimetics (figure 3). This finding is consistent with an upregulation of the p53 family and OPA1 in venetoclax resistant AML cells, and was unexpected due to the roles of the p53 family in inducing pro-apoptotic signalling pathways (41,54).

The role of TAp73 in regulating mitochondrial dynamics may also extend to influencing broader aspects of cellular physiology and proliferation, as highlighted by the fact that following the ablation of TAp73, the growth rates of TAp73 KO cells were significantly impaired (figure S2c). Indeed, the positive regulation of OPA1 is broadly indicative of a growth promoting role of TAp73 isoforms, and should be addressed in further studies. This pro-growth function is consistent with a number of other roles of TAp73, such as the recently reported observation that it is required for cell cycle progression in H1299 cells (55), the regulation of glycolytic flux (15), and the regulation of ETC complexes (16). Overall, these functions contrast with the over- simplified tumour suppressor dogma regarding TA-containing isoforms, and could explain why TAp73 expression is upregulated in many human cancers. Such a phenomenon may represent an out of context activation of the developmental roles of p73 (12).

In addition, we have demonstrated that OPA1 expression is downregulated in the ciliated tracheal epithelium of *Trp73^-/-^*mice, revealing a relevance for the TAp73/OPA1 axis *in vivo* (figure 4). In line with this observation, alterations in mitochondrial morphology and ultrastructure were evident in the *Trp73^-/-^* airway epithelium (figure 4; (9)). Moreover, the downregulation of OPA1 was confined to the ciliated cell lineage, specifically implicating TAp73 in modulating its expression (figure 4). Consequently, we propose that the disruption of mitochondrial function in ciliated epithelial cells may drive the phenotype of dysfunctional ciliogenesis, thereby causing COPD pathogenesis. Moreover, previous studies have shown that the pathogenesis of COPD is caused by alterations in mitochondrial dynamics (56). However, functional assays rescuing MCC differentiation and function with OPA1 expression in the p73 null background are necessary to show that this axis is critical for multiciliated cell function.

It is also notable that *OPA1* heterozygous mice display abnormal brain development, thereby recapitulating features of neurological impairment observed following p73 loss (57,58). Inactivating *OPA1* mutations have been associated with human neurological disorders (59,60), and is driven by an altered transcriptional circuitry in neural stem cells (61). Given this striking similarity with the role of p73 as an essential regulator of neural stem cell maintenance in embryonic and adult neurogenesis (57,62), future work may uncover a molecular relationship between TAp73 and OPA1 in neuronal cells or development of the nervous system. For example, the mechanism underpinning the loss of Cajal-Retzius neurons in the developing mouse hippocampus following oblation of TAp73α remains to be determined (10).

In summary, we have elucidated a novel transcriptional role for TAp73 in regulating mitochondrial dynamics. The regulation of OPA1 adds to the repertoire of p73 functions that are independent of the DNA-damage response. An understanding of such molecular mechanisms is important for paving the way for targeting p73 in conditions such as COPD. In addition to pathological conditions evident from p73 knockout mouse models, a relationship has been defined between cigarette smoke, decreased p73 expression and epithelial cell differentiation *in vivo* (63). Taken alongside the role of TAp73 in regulating important ciliogenesis genes (20), our findings show that alterations in the p73/OPA1 axis may represent a mechanistic link between MCC loss, chronic inflammation and airway disease (figure 5,6). Furthermore, TAp73-dependent regulation of mitochondrial morphology may also play a role in brain development and tumorigenesis.

## METHODS

### Cell Culture & Transfection

NCI-H1299 cells (ATCC #CRL-5803) were cultured in modified RPMI 1640 (Gibco #A4736401, ThermoFisher, Waltham, MA, USA) containing 10% FBS (Gibco) at 37°C and 5% CO_2_.

For transfection, cells were seeded at a density of 400,000 cells per well in 6cm dishes and incubated overnight before transfection with Lipofectamine 2000 reagent (Invitrogen). Lipid-DNA complexes were formed in Opti-MEM medium (Gibco) before addition to cells. The media was replaced after 6h and cells incubated for a further 24-42h depending on the downstream application. Expression plasmid for OPA1 was purchased from Origene (#SC128155, Rockville, MD, USA), and HA- TAp73 cloned into the pcDNA3 backbone.

### TAp73 knockout generation using CRISPR/Cas9

H1299 cells were seeded at a density of 500,000 cells per 6cm dish in 5 mL growth medium and incubated for 24 hours. Cells were transfected with 2 μg Cas9 expression plasmid (Horizon Discovery, Cambridge, UK) and 10 μL gRNA stock (2 μM; Custom Synthesis, Horizon Discovery; CAGGTGGAAGACGTCCATGCT). Transfection mixtures were prepared in 500 mL OPTI-MEM (Gibco) containing 5ul lipofectamine before addition to cells. Cell culture media was replaced after 6h and cells incubated for a further 18 hours. Puromycin was added at a final concentration of 2.5 μg/μL and incubated for 16 hours. Clonal cell populations were obtained by dilution plating in 96-well plates. Once expanded, candidate clones were screened by western blot for TAp73.

Exon 2 spanning PCR fragments were amplified from gDNA of clonal CRISPR populations and cloned into the pJET1.2 blunt cloning vector using the CloneJET PCR cloning kit (Thermofisher, Waltham MA, USA). The ligation reaction was performed following the manufacturer’s instructions and using 1 μL of PCR product. NEB 5-alpha competent E.coli (NEB #C2987H, Ipswich, MA, USA) were transformed with ligated plasmid by heat shock at 42°C for 30 seconds. Cells were spread on pre-warmed LB Agar plates containing 100 μg/mL ampicillin (Sigma, St. Louis, MO, USA) and the plates incubated overnight at 37°C. A minimum of 6 colonies were picked for overnight culture at 37°C, shaking at 230 rpm. Plasmid DNA was obtained using the QIAprep Spin Miniprep Kit (Qiagen #27106, Hilden, Germany). Sanger sequencing was performed by SourceBioscience (Cambridge, UK). Sequencing reads were aligned to the human TAp73 gene (GRCh38) using Geneious Prime software and INDELS identified.

### Animals

*Trp73^-/-^* mice were previously generated in the BALB/c background (11). All procedures followed guidelines and legislation as regulated under the Animal’s Scientific Procedures Act 1986 (ASPA) and were approved by the University of Leicester Animal Welfare and Ethical Review Body (AWERB). For all experiments excluding electron microscopy (EM), mice were euthanised by CO_2_ asphyxiation to preserve the trachea prior to dissection. For EM experiments, anaesthetised alexander mice were perfused via transcardiac perfusion.

### RT-qPCR

RNA was extracted from cell pellets using the RNeasy Plus mini kit (Qiagen #74136) according to the manufacturer’s instructions. For RT-qPCR of epithelial cells, dissected mouse tracheas were transferred to a 10cm cell culture dish containing Ham’s F12 (Gibco) on ice. The trachea were cut open longitudinally to expose the inside surface. Three trachea from either wild-type or *Trp73^-/-^* mice were pooled together in 15 mL falcon tubes containing 2 mL 0.15% filter sterilised Pronase (Roche, Basel, Switzerland) and incubated at 4°C overnight. The tubes were gently inverted and FBS (Gibco) added at a final concentration of 10%. The tracheas were then transferred to a new 15ml tube containing Ham’s F12/10% FBS and inverted. The contents from each tube were pooled and cells centrifuged at 300g, 4°C for 10 mins. The pellet resuspended in 600 μL DNase solution (Sigma; 0.5 mg/mL crude pancreatic DNase I, 10 mg/ml BSA, in Ham’s F12), and incubated on ice for 5 mins. RNA was extracted using TRIzol reagent (Invitrogen), according to the manufacturer’s instructions.

Reverse transcription reaction was performed using RevertAid minus first strand cDNA synthesis kit (Thermofisher). Transcribed cDNA was diluted 2x before use in qPCR. Primer sequences (Sigma) are listed in Table 1. Reactions were carried out in triplicate using Fast SYBR Green PCR Master Mix (Thermofisher #4385612). The relative quantification was obtained using the Applied Biosystems 7500 thermocycler and quantitative comparative (ΔΔCt) method normalised to TBP.

**Table 1.**
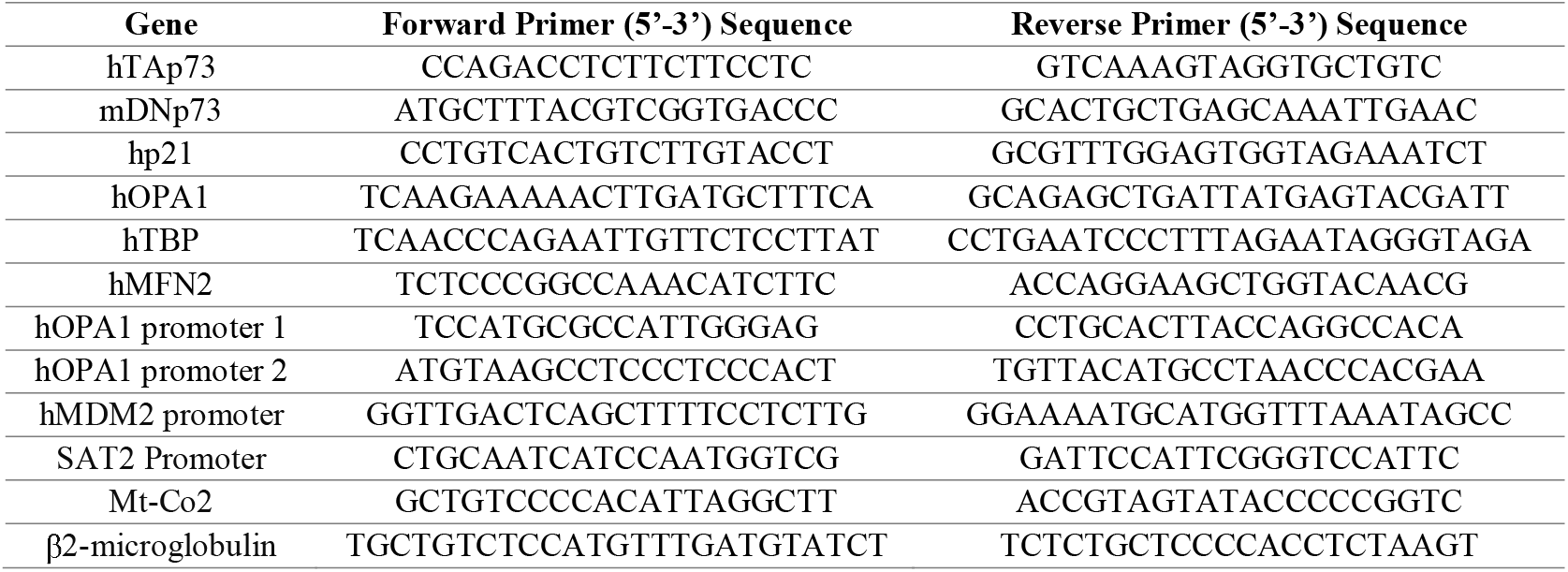
A list of primer sequences used for qPCR.

### Chromatin Immunoprecipitation

The MAGnify™ chromatin immunoprecipitation kit (Invitrogen) was used according to the manufacturer’s instructions. H1299 cells were transfected with TAp73α overexpression construct as described previously. Cells were fixed by adding PFA at a final concentration of 1% and incubating for 10 minutes at RT. Cells were lysed according to the manufacturer’s instructions and stored at -80°C. Chromatin was sheared using the Covaris S220 set at a duty factor of 2%, intensity 4, and 200 cycles/burst for 6 mins. Anti-HA antibody or mouse immunoglobulins was bound to Dynabeads® by rotating end-over-end at 4°C for 1h prior to IP. Chromatin was diluted to 200,000 cells per IP and immunoprecipitated according to the manufacturer’s instructions and eluted in 150 μL elution buffer.

### Western Blot

Cell pellets were lysed using RIPA buffer (Merck, Darmstadt, Germany) supplemented with 0.1 % (v/v) protease and phosphatase inhibitor cocktail (Sigma-Aldrich, US). Protein concentration was measured using the Bio-Rad protein assay.

Proteins were resolved by SDS-PAGE using 4-20 % or 4-15% TRIS-glycine gradient midi gels (Bio-Rad, Hercules, CA, USA), in electrode buffer (0.1% w/v SDS, 192 mM glycine, 25 mM Tris; Bio-Rad) in BioRad criterion tanks. Protein was then transferred overnight onto a nitrocellulose membrane (Immobilon-P, Merk) using wet transfer for 16 h at 25V in transfer buffer (25 mM Tris, 192 mM glycine and 20 % methanol). The membranes were then blocked with 5% non-fat dry milk (Marvel) in TBST (0.1%) for 1h at room temperature. The membrane was then probed with the appropriate primary antibody (Table 2) for 1h at room temperature and washed with TBST. Membranes were then incubated with HRP conjugated secondary antibodies for 1h at room temperature. Clarity Max Western ECL substrate was added to the membranes for 5 mins (Bio-Rad #1705062) and chemiluminescent signal imaged using a Chemidoc imager (Bio-Rad).

**Table 2.** A list of antibodies used in Western Blotting experiments.

| Antibody Target | Manufacturer | Catalogue Number | Host Species | Antibody Dilution |
| --- | --- | --- | --- | --- |
| TAp73 | Bethyl | A300-126A | Rabbit | 1:1000 |
| OPA1 | BD Biosciences | 612606 | Mouse | 1:1000 |
| Mitofusin-1 | Cell Signaling | 14739 | Rabbit | 1:1000 |
| Mitofusin-2 | Cell Signaling | 9482 | Rabbit | 1:1000 |
| p21 | Santa Cruz | Sc-6246 | Mouse | 1:200 |
| ATP5B | Abcam | Ab14730 | Mouse | 1:1000 |
| GAPDH | Sigma-Aldrich | G8795 | Mouse | 1:5000 |
| HSP-90 | Santa-Cruz | Sc-13119 | Mouse | 1:1000 |
| Total Human OXPHOS<br>Cocktail | Abcam | Ab110411 | Mouse | 1:1000 |
| MCL-1 | Rockland | 600-401-394 | Rabbit | 1:1000 |
| BCL2 | Santa-Cruz | Sc-509 | Mouse | 1:200 |
| BCL-xL | Genetex | GTX100064 | Rabbit | 1:1000 |
| Vinculin | Abcam | Ab18058 | Mouse | 1:5000 |

### Immunofluorescence

H1299 cells were seeded onto sterile glass coverslips (Type 1.5 thickness) and cultured overnight. Cells were fixed in 4% paraformaldehyde (VWR, Radnor, PA, USA) at room temperature for 20 min. The cells were washed 3 times in PBS and permeabilised at room temperature for 20 min in 0.2% Triton-X 100/PBS. Cells were washed with PBS/Tween (0.1%) 3 times, cover slips transferred to 12-well plates, and blocked in 5% Goat Serum/PBS/Tween for 1h at room temperature. Anti-ATP5B antibody (mouse monoclonal, Abcam #ab14730, Cambridge, UK) was diluted 1:1000 in PBS and cover slips incubated overnight at 4°C. Cells were washed 3 times with PBS/Tween (0.1 %) and incubated with Alexa Fluor 488 goat anti-mouse secondary antibody diluted 1:1000 in PBS for 1 h at room temperature whilst protected from light. Cells transfected with HA-TAp73-pcDNA3 plasmid were incubated with rabbit anti-HA antibody (Santa Cruz #805, Dallas, TX, USA) diluted 1:200 in PBS for 1h at room temperature, followed by detection with Alexa Fluor 568 goat anti-rabbit secondary antibody diluted 1:1000 in PBS. Coverslips were mounted on glass slides with Vectashield plus antifade mounting medium with DAPI (Vector Laboratories #H-2000, Newark, CA, USA). Images were acquired using a Zeiss LSM880 confocal microscope and 63x oil objective.

### Immunohistochemistry

IHC was carried out using the Ventana Discovery Ultra platform (Roche). Deparaffinization was performed by immersion in Ventana Discovery Wash buffer (Roche; 950-510) for 3 x 8 minute washes at 69°C. Antigen Retrieval was performed by incubation in Ventana CC1 buffer at 95°C for 32 minutes. The slides were blocked by incubation with Discovery goat IgG block (Roche; 760-6008) for 12 mins at 37°C. Antibodies were diluted in EnVision Flex Antibody Diluent (Agilent; DM830, Santa Clara, CA, USA), at the dilutions indicated in table 2. Multiplex antibody detection was performed using Opal fluorophores according to the manufacturer’s instructions (OPA1; Opal 690, Ac-α-tubulin; Opal 540; FOXJ1; Opal 570). Slides were imaged using the Ventana Discovery imaging system and quantification of signal intensity performed using In-form software.

### Incucyte Annexin V/APC imaging

H1299 cells were seeded at a density of 100,000 cells/well in a 12 well plate and cultured overnight at 37°C and 5% CO_2_. Cell culture media was aspirated and fresh media added containing 1:3000 Annexin V/APC dye (made in house by Dr Xiao- Ming Sun), 1 μM CaCl_2_, and incubated for a further 30 mins. BH3-mimetics ABT- 737 (Inhibitor of BCL-2 and BCL-xL) and S63845 (Inhibitor of MCL-1) were added to cells at a range of concentrations ranging from 0.25um-10 μm per well or vehicle control (0.1% DMSO) and the cell culture plate placed in the IncuCyte live cell imager. Assay plates were scanned using the adherent cell-by-cell module and the 20x objective. Label-free counts were obtained using the phase contrast channel with a segmentation adjustment of 0.2 and a minimum area of 50 μm^2^. The green channel was imaged using an acquisition time of 300 ms and objects counted using the Top- Hat segmentation method with a threshold of 2.0 green calibrated units.

### Seahorse Extracellular Flux Assay

Cells were cultured overnight at 37°C, 5% CO_2_ in 96-well Seahorse microplates. Growth media was then removed, and cells washed three times in unbuffered DMEM Seahorse assay medium (32 mM NaCl, 2 mM GlutaMAX, 1 mM sodium pyruvate, 11 mM D-glucose, pH 7.4). The plate was then incubated in a 37 °C non-CO_2_ incubator for 1h. The canonical mitochondrial toxins oligomycin A (port A), FCCP (port B), antimycin A and rotenone (port C) were added at the following final concentrations; 2 μM, 500 nM, and 2 μM respectively, at 18 minute intervals. Mean OCR values were obtained from a minimum of 5 wells per treatment and background subtracted. The data was normalised to cell number using Hoechst 33342 staining and individual wells were imaged using the 5x objective on a Zeiss Observer 7 microscope fitted with a Colibri 7 LED fluorescence light source.

### Conventional and Large-Volume Electron Microscopy

Mouse tissue sections and cells were fixed with half Karnovsky fixative as 2.5% glutaraldehyde and 2% paraformaldehyde in NaHCa buffer (0.1 dM NaCl, 30 mM HEPES, 2 mM CaCl2, pH 7.4) for 4 hours at room temperature. For conventional transmission electron microscopy (C-TEM)(64), post-fixation was performed with 0.25% osmium tetroxide + 0.25% potassium ferrocyanide and 1% tannic acid in 0.1 M sodium cacodylate buffer (pH 7.4). After staining *en bloc* with 5% aqueous uranyl acetate solution, the dehydration with a series of ethanol and the resin infiltration were completed for the plastic embedding in TER (TAAB Epoxy Resin). Ultrathin-sections (∼60 nm) were cut using an ultramicrotome (Leica EM UCT/UC7/Artos-3D, Austria), mounted in EM grids, and stained with lead citrate. Targets were observed using FEI Talos F200C (ThermoFisher) with Ceta-16M CMOS-based camera (4kx4k pixels under 16bit dynamic range) and JEM-1400 Flash TMP (JEOL Ltd., Japan) with TVIPS TemCam-XF416 CMOS (Tietz Video and Image Processing Systems GmbH, Germany) as described previously (9,10)

For large-volume serial-block-face scanning electron microscopy (SBF-SEM) (65), the tissues and cells were fixed with half Karnovsky’s fixative and post-fixed with osmium-thiocarbohydrazide-osmium (OTO) (1% osmium tetroxide + 1% potassium ferrocyanide in cacodylate buffer, aqueous 1% thiocarbohydrazide solution and aqueous 2% osmium tetroxide solution). *en-bloc* staining of fixed tissues was performed with aqueous 5% uranyl acetate solution. After dehydration and infiltration, samples were embedded in epoxy TER resin. Target regions were trimmed-down, mounted on aluminium pin stubs with conductive epoxy (CircuitWorks CW2400, Waukegan, IL, USA) and imaged by SBF-SEM in FEI Quanta FEG 250 (Thermo Fischer Scientific) equipped with ‘3View2XP’ system (Gatan Inc, Pleasanton, CA, USA) as described previously (66). Back-scatter- electrons from the serial block-faces were recorded by Gatan ‘OnPoint’ detector at a pixel size 2.5 nm and beam acceleration of 3 kV under lower vacuum state of 40 Pascal, using the following parameters: spot size of 3.5, a dwell-time of 1.5 μs per pixel, ROI size of 10,000 x 25,000 (XY) pixels, 3 ROIs acquired for the montage, Z-slice thickness of 80 nm. 3D segmentation and reconstruction for the cellular tissue organelles were done by FEI Amira-3D, IMOD-4 (67), EMAN-2 (68).

### Human Lung Genomic Data

Genomic data was provided by the Lung Genomics Research Consortium (LGRC; http://lung-gemomics.org; 1RC2HL101715) using tissue samples and clinical data collected through the Lung Tissue Research Consortium (LTRC; http://www.ltrcpublic.com/), data series GSE47460. Gene expression analysis was performed using the PulmonDB tool (69).

### ChIP-seq Analysis

Publicly available ChIP-seq data sets (GEO series GSE15780) against p53 and p73 isoforms (TAp73α and TAp73β) in the human osteosarcoma cell line Saos-2 were interrogated for genes associated with the regulation of mitochondrial fusion and fission (70). BED files were converted to WIG files, and UCSC Genome Browser annotated tracks generated using the ChIP-Seq tools and web server (71). ChIP-seq tracks were generated for each replicate and cross-checked against input tracks to account for non-specific signal. The maximum Y-axis scale was set to 100 for each genomic loci to a allow for relative depiction of signal strength.

### Gene Ontology Analysis

GO analysis was performed on the complete set of genomic binding sites of p73 (1769 genes) identified *In-Situ* by Marshall et. al. (20) using dissected murine trachea. The list of genes was analysed using the PANTHER GO tool v.17 overrepresentation test (72). Genes were analysed against the complete set of Mus musculus genes annotated with the GO biological process complete annotation data set. Fold-enrichment values were plotted for selected GO classes (p = <0.01).

## ACKNOWLEDGEMENTS

We sincerely thank Catherine Ficken and Mark Southwood for the extensive technical support with immunohistochemical staining and analysis. We also thank Maria Guerra Martin for support with EM sample processing.

This work was funded by the UK Medical Research Council, intramural project MC_UU_00025/4 (RG94521) to M.M.F. and MC_UU_00025/2 to G.M. The funders had no role in study design, data collection and analysis, decision to publish or preparation of the manuscript.

## AUTHOR CONTRIBUIONS

G.M. and M.M.F. initiated the project. N.B, M.M.F., L.M.M., G.M., I.A., E.P., L.P., and A.C. designed the study, coordinated the experiments, and provided conceptual inputs for the paper. N.B. wrote the manuscript with input from M.M.F. and L.M.M. N.B., X.M.S., N.M. and J.L. performed the experiments and analysed the data. All authors read and approved the final manuscript.

## SUPPLEMENTARY FIGURES

**Figure S1.**
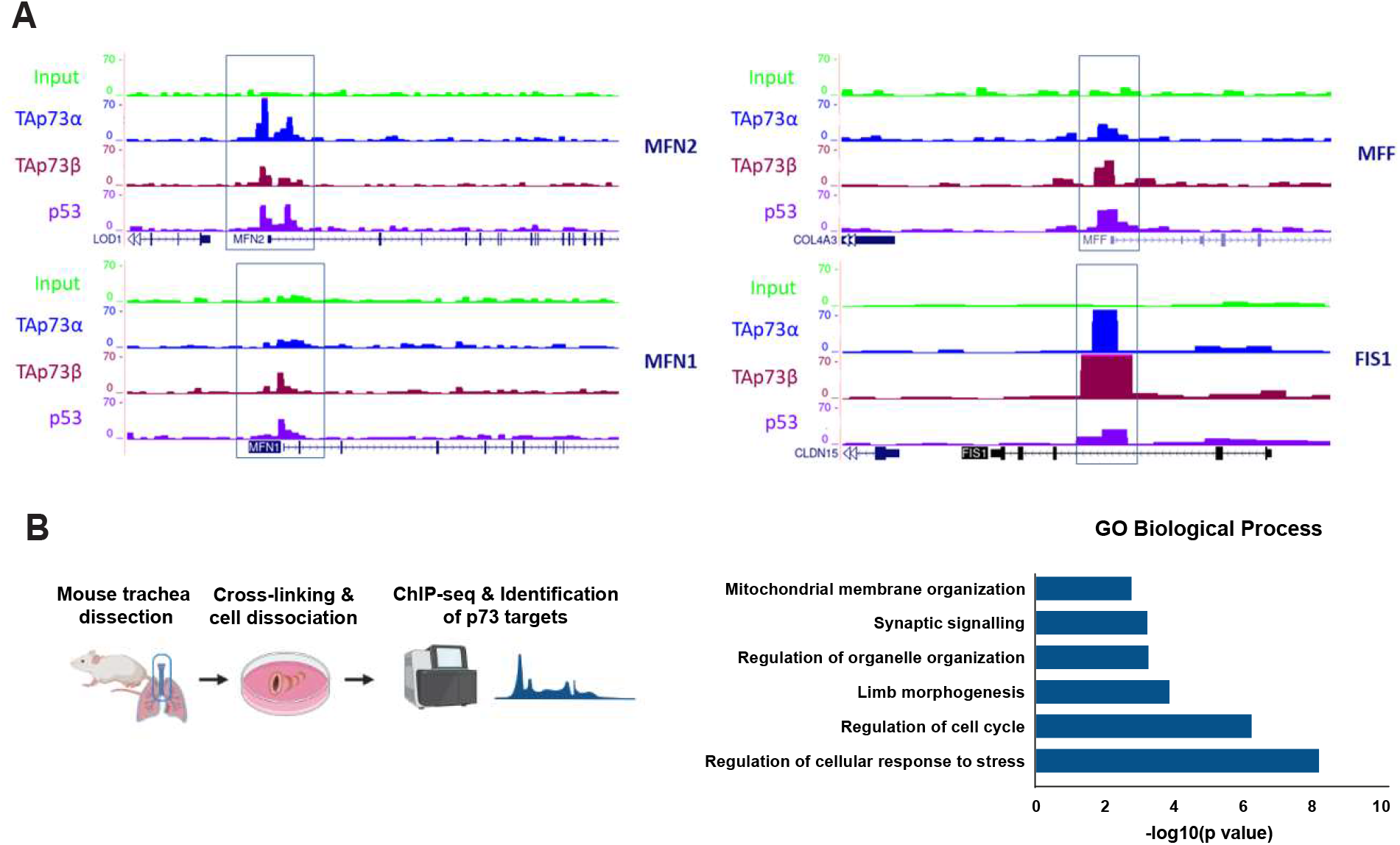
ChIP-seq data shows TAp73 and p53 binding to a range of genes that regulate mitochondrial morphology. (A) Interrogation of ChIP-seq data indicated binding of TAp73α, TAp73β and p53 to the *MFN2*, *MFF* and *FIS1* gene loci. Input signal is shown in green. Sequencing read files were obtained from the GEO data set GSE15780. (B) GO enrichment analysis of 1769 TAp73 target genes identified using *in-situ* mouse tracheal ChIP-seq. Analysis was performed using the Panther GO overrepresentation test.

**Figure S2.**
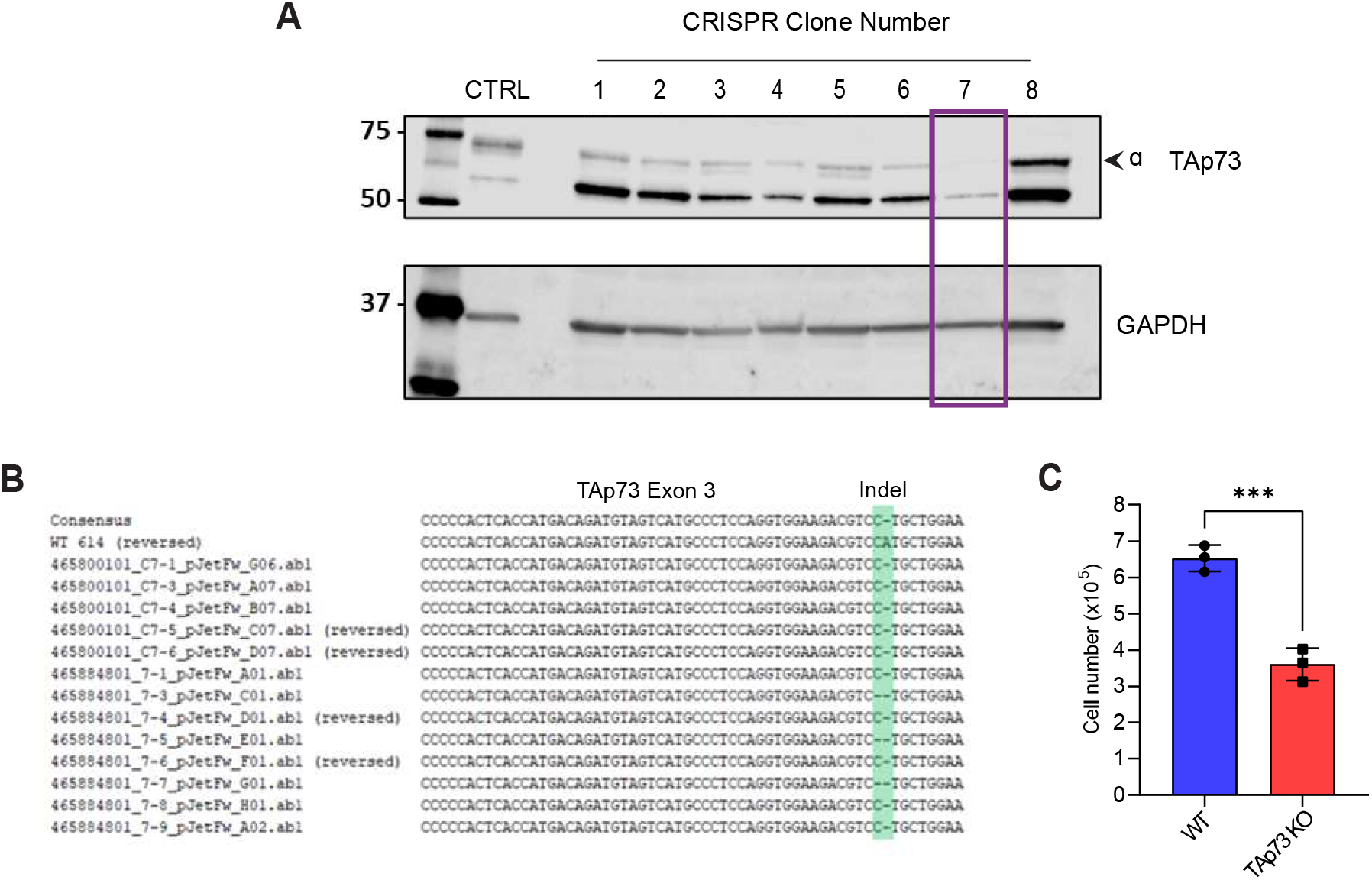
Generation of TAp73 KO cell lines using CRISPR/Cas9 targeting. (A) Representative western blot carried out to screen for TAp73 KO H1299 clonal populations. Clone 7 (purple box) was selected for Sanger Sequencing. (B) Sanger sequencing reads obtained from TAp73 KO CRISPR clone 7 aligned to the WT sequence. Primers were designed to amplify exon 3 of the *Trp73* gene, encompassing the target site. The site of gene editing (INDEL) is shown in the green box at the DNA bases where CRISPR cells had a single or double base pair deletion immediately upstream of the PAM sequence. (C) Cells from the indicated cell lines were seeded at 20,000 cells/well in a 6 well plate and counted after 5 days to obtain relative rates of proliferation. Data are shown as mean ± SD (n=3). (***) P ≤ 0.001 (Student’s T-test).

**Figure S3.**
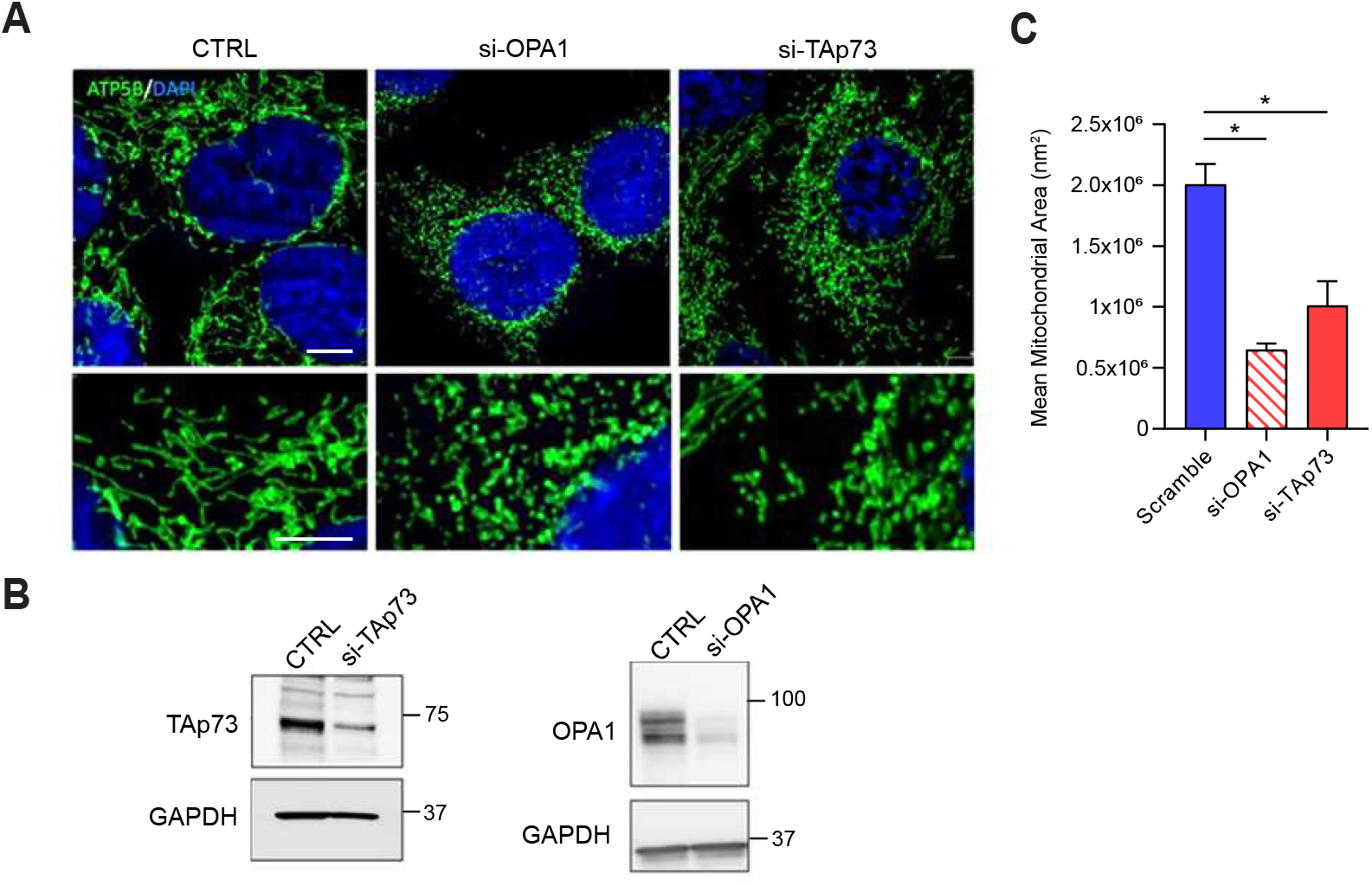
H1299 cells treated with siRNA targeting TAp73 display fragmentation of the mitochondrial network. (A) Knockdown of TAp73 was induced by siRNA. Cells were subsequently fixed and probed for ATP5B to visualise the mitochondrial network by immunofluorescence (green). The observed phenotype in TAp73 knockout cells is consistent with a depletion of OPA1 (middle panel). (B) siRNA mediated knockdown of TAp73 and OPA1 was confirmed by western blot. Images representative of two independent experiments (n=2). (C) Quantification of mitochondrial area in WT and si-TAp73 treated cells using Intellesis trainable object segmentation. (*) P ≤ 0.05 (Student’s T-test).

**Figure S4.**
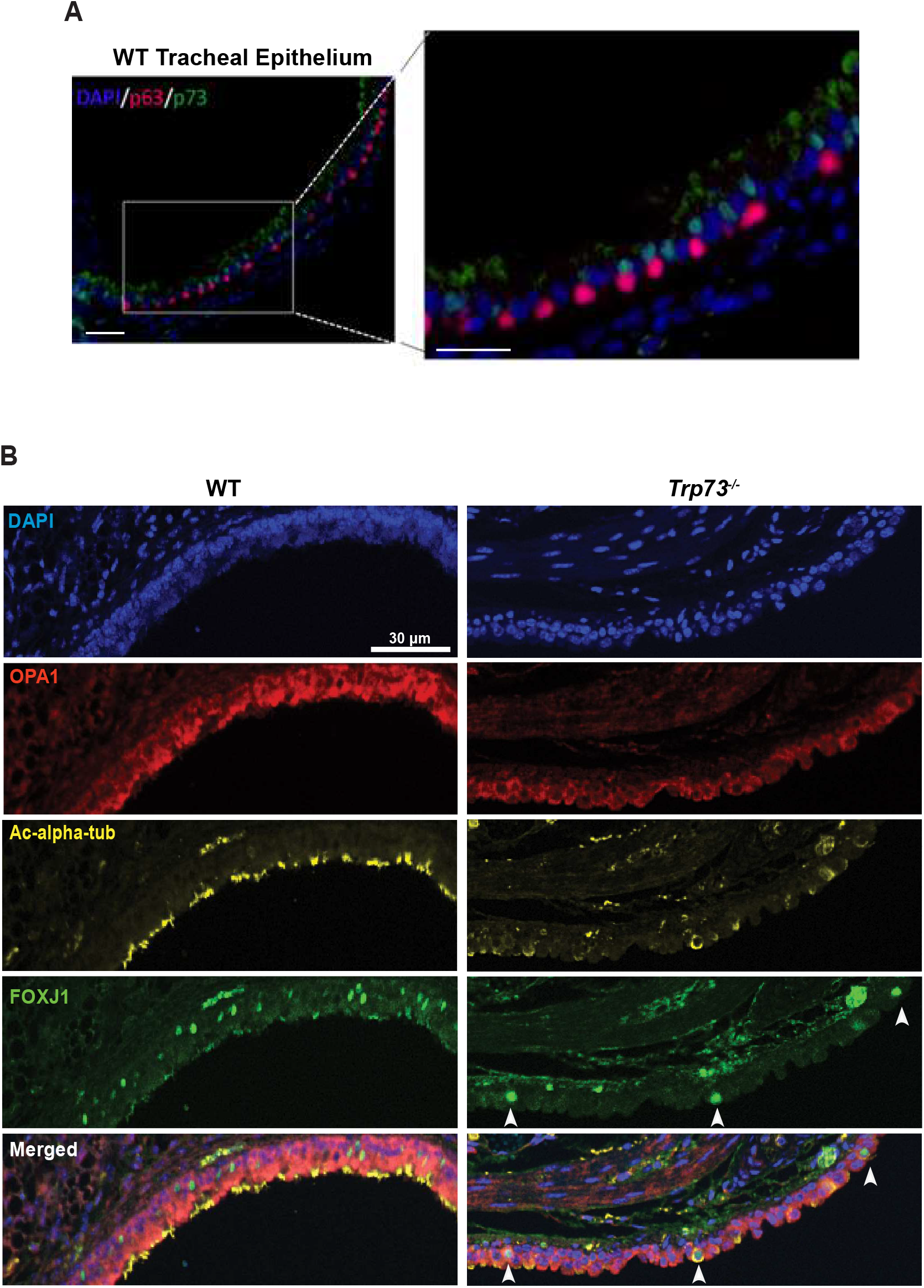
TAp73 is not expressed in the basal cell compartment of the mouse tracheal epithelium, and the *Trp73^-/-^* epithelium maintains FOXJ1 expression. (A) Dual immunohistochemical staining for TAp73 (green) and p63 (pink) in the wild-type tracheal epithelium. No co-localisation of the signal was observed. (B) Single channel images from multiplex IHC staining performed in figure 4. Staining for FOXJ1 was also incorporated. White arrow heads indicates FOXJ1 positive cell populations with perturbations in cilia structure.

**Figure S5.**
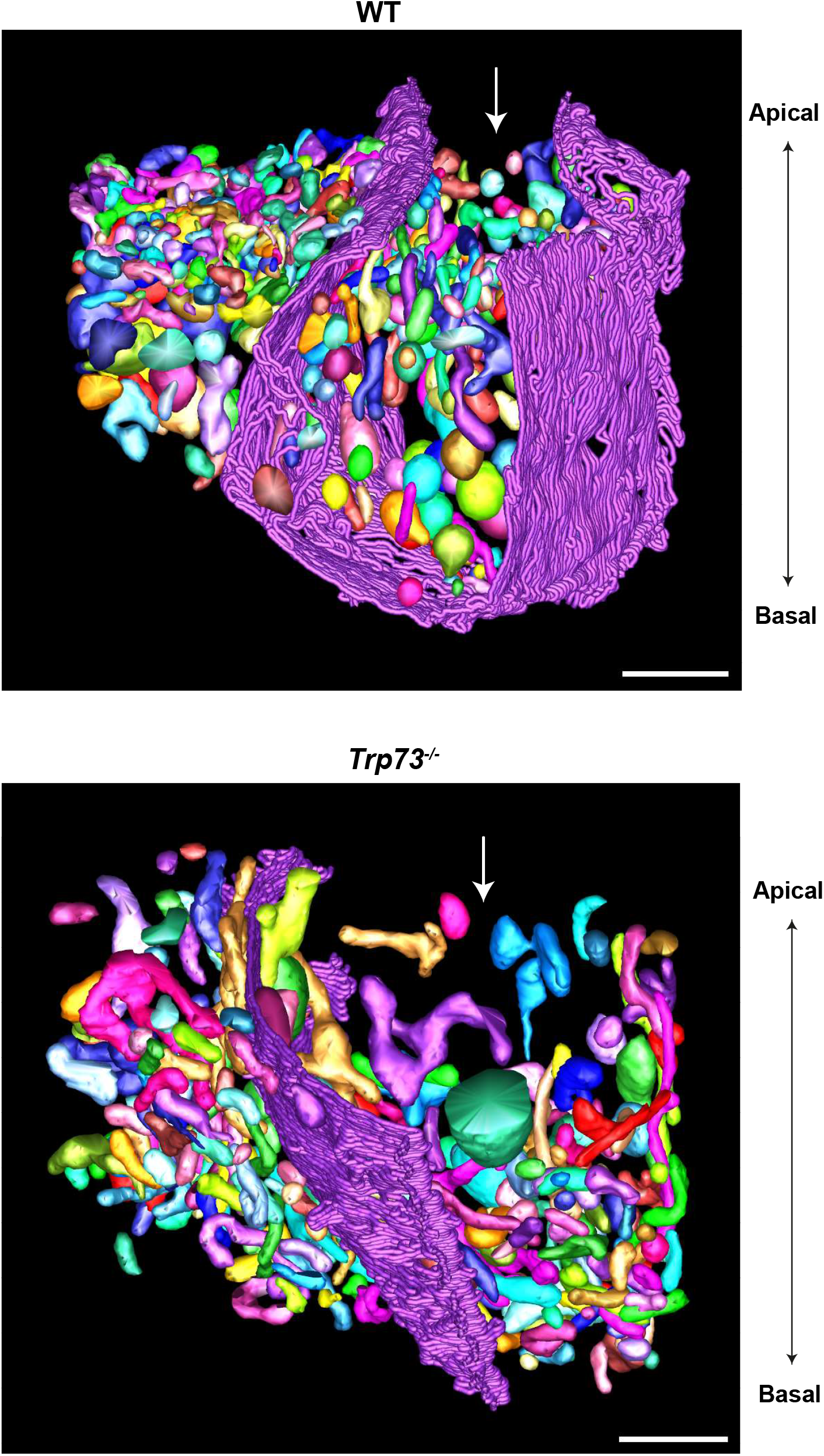
SBF-SEM reconstruction of the mitochondrial network in ciliated cells of the tracheal epithelium. Mitochondria were segmented from 100 sequential sections of tracheal epithelial cells (WT and *Trp73^-/-^*) to construct a 3D model of the mitochondrial network. White arrows denote ciliated epithelial cells, with cell boundaries indicated in purple. The apical and basal surfaces are indicated. Scale bar = 2500 nm.

